# Clustering temporal step-counting patterns for 24 hours with machine learning revealed potential heterogeneity in the categorization by a traditional tertile procedure

**DOI:** 10.1101/2023.06.04.543652

**Authors:** Saida Salima Nawrin, Hitoshi Inada, Haruki Momma, Ryoichi nagatomi

**Author notes:** Corresponding author: (HI), (RN). Department of Biochemistry & Cellular Biology, National Center of Neurology and Psychiatry, Kodaira, Tokyo, Japan (Hitoshi Inada). These authors contributed equally to this work (Saida Salima Nawrin, Hitoshi Inada).

## Abstract

**Objectives:** Physical activity is a crucial aspect of health benefits in the public society. Although studies on the temporal physical activity patterns might lead to the protocol for efficient intervention/program, a standardized procedure to determine and analyze the temporal physical activity patterns remains to be developed. Here, we attempted to develop a procedure to cluster 24-hour patterns of physical activity as step counts measured with an accelerometer-based wearable sensor.

**Methods:** Data was collected from 42 healthy participants, comprising 35 males and 7 females, at the Sendai Oroshisho center in 2008. Using unsupervised machine learning, specifically the kernel k-means algorithm with the global alignment kernel was applied on a total of 815 days from 42 participants, and 6 activity patterns were identified. Further, the probability of each 24-hour step-counting pattern was calculated for every participant., and was used in MATLAB to apply spectral clustering, and 5 activity behaviors were identified.

**Results:** We could identify six 24-hour step-counting patterns and five daily step-behavioral clusters. When the amount of physical activity was categorized into tertile groups reflecting highly active, moderately active, and low active, each tertile group consisted of different proportions of six 24-hour step-counting patterns.

**Conclusions:** Our study introduces a novel approach using an unsupervised machine learning method to categorize daily hourly activity, revealing six distinct step counting patterns and five clusters representing daily step behaviors. Our procedure would be reliable for finding and clustering physical activity patterns/behaviors and reveal heterogeneity in the categorization by a traditional tertile procedure using total step amount.

## Introduction

Physical activity is a crucial aspect of maintaining good health and well-being. World Health Organization (WHO) defines “physical activity as any bodily movement produced by skeletal muscles that requires energy expenditure” and recommends at least 75-150/150-300 minutes of vigorous/moderate-intensity aerobic physical activity throughout the week (Organization WHO. Physical Activity 2022). WHO also mentioned that physical inactivity is one of the leading risk factors for global mortality, causing an estimated 3.2 million deaths per year (Organization WHO Physical Inactivity). Many observational and intervention studies suggested that regular physical activity has been linked to numerous physical and mental health benefits, including reducing the risk of chronic diseases, improving cardiovascular health, maintaining a healthy weight, improving mental health, and enhancing the overall quality of life (Chekroud et al., 2018; Kraus et al., 2019; Madigan et al., 2021; Pearce et al., 2022; Piercy et al., 2018). However, a recent systematic review and meta-analysis of randomized controlled trials indicated that interventions to maintain physical activity behavior was effective in short periods (at least three months) but provided a small impact on the long-term maintenance of physical activity (Madigan et al., 2021). Therefore, it is critical to figure out efficient ways integrated into daily life to promote and maintain physical activity for the improvement of the health and well-being of the public.

A historical topic in public health is what kind, how much, and how intense physical activity is required for health benefits (Powell, Paluch, & Blair, 2011). One of the simplest ways to increase and maintain physical activity is to walk/move more and longer (Morris & Hardman, 1997; National Heart, Lung, and Blood Institute. Tips for Getting Active; Ogilvie et al., 2007) as shown in the 10,000-steps campaign (Eyler, Brownson, Bacak, & Housemann, 2003; Hatano, Y. 1993; Tudor-Locke & Bassett, 2004). A recent meta-analysis reported that taking more steps per day was associated with lower mortality risk, where the risk plateaued at approximately 6,000–8,000 steps or 8,000–10,000 steps per day for older (≥60 years) or younger adults (<60 years), respectively (Paluch et al., 2022). Most of the previous research on physical activity has concentrated on how the total amount of activity based on its intensity and duration is linked to health (Amagasa et al., 2018; Brailey et al., 2022; Lu et al., 2022). However, analyzing only the total amount of physical activity might overlook a complete understanding of the activity behavior since the metrics of the total amount of physical activity tends to exclude or underestimate the effect of temporal variations of physical activity.

Recently, several studies focused on the 24-hour temporal physical activity patterns using a narrower criterion for physical activity, such as steps or body movements objectively measured with an accelerometer (Aqeel et al., 2021; Niemelä et al., 2019; Smagula et al., 2015; Smagula, Krafty, Thayer, Buysse, & Hall, 2018; Smagula et al., 2022). An earlier study that analyzed the relationship between activity rhythm disturbances and depression identified eight sub-clusters in older adults (≥65 years) (Smagula et al., 2015). Late or combined early/compressed/dampened activity rhythms may independently contribute to depression symptom development (Smagula et al., 2015). Another recent report using activity counts for seven days in US adults indicated that, although the lowest physical activity counts cluster showed a higher obesity index including body mass index and waist circumference than the other clusters showing higher activity counts, the clusters also expressed different temporal patterns of physical activity (Aqeel et al., 2021). Moreover, an increase in cardiovascular disease risk was observed in the inactive cluster than the other clusters (evening active, moderately active, and very active) (Niemelä et al., 2019) from the data of The Northern Finland Birth Cohort 1966 study (Northern Finland Birth Cohorts | University of Oulu). The latest report combining activity timing/period and pattern robustness identified sub-groups with progressive depression symptoms and impaired cognitive performance in aging (Smagula et al., 2022). Therefore, understanding temporal physical activity patterns would help extract informative health outcomes.

In the previous studies above, participants were first clustered by several selected features indicating physical activity patterns, and the physical activity patterns in 24 hours were analyzed. However, those procedures have difficulty analyzing temporal physical activity patterns directly because of their methodological restrictions, in which participants were first clustered by extracted individual features of the physical activity patterns and then analyzed. In this study, we propose a novel procedure for the direct clustering of physical activity based on the temporal step-counting patterns with unsupervised machine learning irrespective of the total volume of physical activity. Subsequently, we could identify five step-behavioral clusters of participants, based on the proportion of each step-counting pattern. Furthermore, we analyzed the proportion of the temporal step-counting patterns and step-behavioral clusters of groups categorized by a traditional tertile procedure The results revealed potential heterogeneity in categorizing by the tertile procedure using total step amount. Our approach could provide a useful procedure to analyze temporal patterns of physical activity in a direct manner.

## Materials and Methods

### Participants and data collection

We used the anonymized activity counting data obtained in the previous study (Guo et al., 2012), in which participatns were recruited in August 2008 throughout an annual health examination at the Sendai Oroshisho center, provided and agreed with informed consent for their data to be analyzed (Guo et al., 2012). The data collected from forty-two participants between August and September 2008 in Sendai Oroshisho center was used for a series of analyses. The Institutional Review Board of the Tohoku University Graduate School of Medicine have approved the current study protocol (Approval number: 2019-1-394).

Step count data was continuously collected by a hip-worn triaxial accelerometer (Fig S1, Nipro Welsupport, model no wat3663, Japan) for up to 25 days. Data from participants with less than three days was excluded for statistical reasons (Hart, Swartz, Cashin, & Strath, 2011). The demography of the forty-two participants (35 males and 7 females). Each participant has 19 ± 4 days of data. The data sampling rate was 1 Hz. Eight types of activities are recorded as below: 0, Resting; 1, Moving while standing; 2, Going down using a vehicle; 3, Going up using vehicle; 4, Going down using stairs; 5, Walking on the ground level; 6, Going up using stairs; 7, Running; 8, Lying down. Numbers of indices 5-7 were counted as steps. The other indices were counted as zero.

### Software for data processing and visualization

MATLAB (2021a, MathWorks, MA) and Python (3.11.0) were used for data processing and visualization.

#### Twenty-four-hour step-counting pattern

The activity counting data at 1 Hz sampling rate was first converted hourly for each 24-hour. A total of 815 days of hourly data was used to cluster the 24-hour step-counting patterns by unsupervised machine learning using tslearn, a Python package for time-series analysis (Tavenard et al., 2020). The kernel k-means algorithm was used for clustering with the global alignment kernel (GAK). The cluster number was determined by elbow plot using distortion values. Six clusters of 24-hour physical activity patterns were obtained.

#### Dominant daily step behavior

First, the probability of each 24-hour step-counting pattern is calculated for each participant based on the formula below.

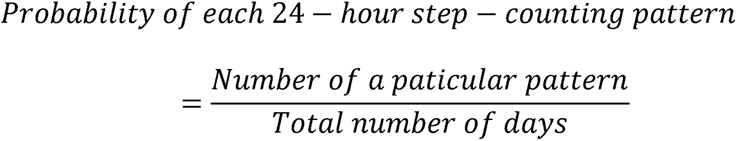

Vectors of the probability of each 24-hour step-counting pattern (with six values) were used to cluster the daily step behaviors by unsupervised machine learning using a spectral clustering algorithm with MATLAB. Five clusters of daily step-behavior were obtained.

### Tertile clustering of daily physical activity based on the daily step amount

The daily step amount for 815 days was divided into the tertile group (low, medium, and high). Each group is defined as:

> Low: Daily step amount <= 4127
>
> Medium: 4127 < Daily step amount < 6946
>
> High: Daily step amount >= 6946

### Statistical analysis

Pearson chi-squared test was performed with (JMP pro 16.0.0) for all statistical analyses where P<.05 is considered significant.

## Results

The data set was collected from forty-two healthy participants (35 males and 7 females) from August to September 2008 in Miyagi prefecture, Japan. The demography of the participants is shown in Table 1.

**Table 1.** Demographical information.

| Variables | Mean ± SD |
| --- | --- |
| Age (Years) | 45.59 ± 10.96 |
| Weight (kg) | 55.83 ± 10.69 |
| Body fat (%) | 29.37 ± 6.29 |
| BMI | 22.10 ± 3.42 |
| Waist (cm) | 78.04 ± 9.72 |

Note: Demographic information for 42 participants

### Twenty-four-hour step-counting pattern

Step count data obtained with a triaxial accelerometer was first converted hourly for each 24-hour and then applied for the clustering with unsupervised machine learning (See Methods). Fig 1 shows six 24-hour step-counting patterns identified: all-day (AD), bi-phasic (BP), morning (M), evening (E), irregular morning (IM), and irregular night (IN). AD showed a continuous high step count from 9:00 to 19:00. BP showed increased step counts from 8:00 to 21:00 but had two peak periods at 9:00 and 19:00. Step counts between 10:00-16:00 was slightly low in BP. M started actively around 6:00, kept on high step counts until 14:00, and gradually decreased in activity after 15:00. By contrast, E started active afternoon and showed peak step counts between 16:00-18:00. IM showed high step counts mainly in the morning from 5:00 to 13:00 and decreases in activity after 15:00. IN showed high step-counts from 19:00 to midnight. Table 2 summarizes the proportion of each step-counting pattern. Among 815 days, AD was the most dominant pattern (330/815, 40.5 %), and the BP pattern followed (146/815, 17.9%). M and E patterns were 15.8 % and 13.0%, respectively. IM and IN were minor patterns, less than 10% (7.9% and 4.9 %, respectively). The proportion of each pattern showed significant differences except for a difference between IM and IN (Table S1). Each step-counting pattern showed a different step-count amount (Table 3). The AD showed the highest step-count amount per day (7020.0 ± 3271.4 counts). The M and BP patterns were similar, and the E pattern followed. The IN and IM patterns showed lower step count amounts. AD, BP, and M patterns differed significantly from E, IM, and IN (Table S2). Among E, IM, and IN patterns, a significant difference was observed between E and IM patterns. Regarding duration, the BP pattern showed the longest duration of activity per day (14.54 ± 2.61 hours). The AD and M followed. Generally, the AD and BP patterns shared 58.4 % of total days and showed higher/longer step count amounts per day. The M pattern was more active than the E pattern. The IM and IN shared a minor proportion in days (12.8 %) and showed the lowest step-count amount. All the pairs show a difference (p < 0.05) except BP-AD (Table S3).

**Fig 1.**
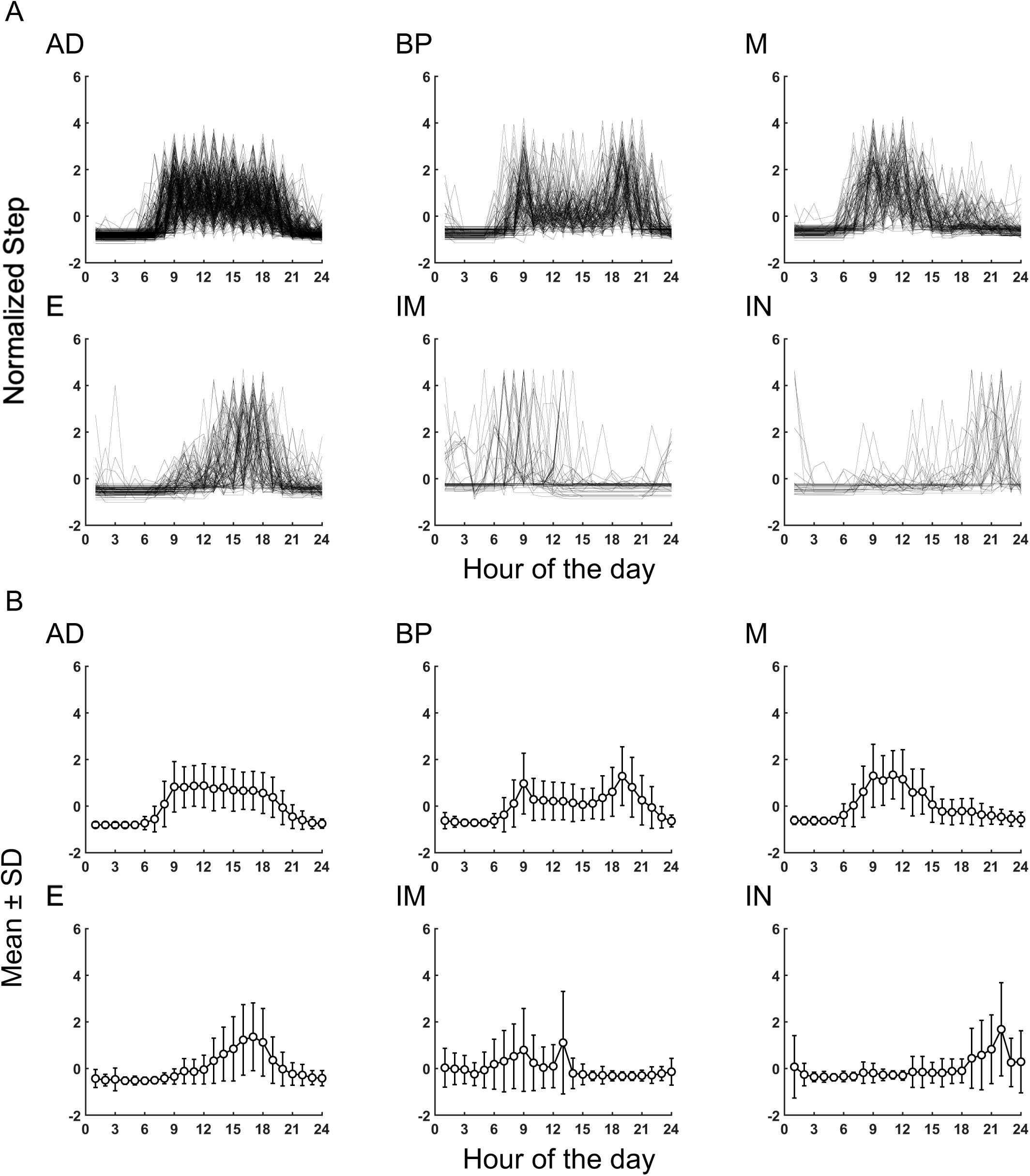
Six step-counting patterns for 24 hours. Temporal step-counting pattern for 24 hours identified using unsupervised machine learning. (A) All traces of step-counting activity in each pattern. (B) Averaged traces (mean ± SD) of step-counting activity in each pattern. AD, all-day; BP, bi-phasic; M, morning; E, evening; IM, irregular morning; IN, irregular night.

**Table 2.** Proportion of each step-counting pattern.

| Pattern | Days | % of days |
| --- | --- | --- |
| All-Day (AD) | 330 | 40.5 |
| Bi-phasic (BP) | 146 | 17.9 |
| Morning (M) | 129 | 15.8 |
| Evening (E) | 106 | 13.0 |
| Irregular morning (IM) | 64 | 7.9 |
| Irregular night (IN) | 40 | 4.9 |
| Total | 815 | 100 |

Note: In total 815 days, the AD pattern was the most common, followed by BP, M, and E. IM and IN patterns were less frequent.

**Table 3.** Step count amount and duration for each pattern.

| Pattern | Step count amount<br>mean ± SD | Activity duration (hours)<br>mean ± SD |
| --- | --- | --- |
| All-Day (AD) | 7020.0 ± 3271.4 | 14.29 ± 2.28 |
| Bi-phasic (BP) | 6178.2 ± 3102.1 | 14.54 ± 2.61 |
| Morning (M) | 6473.7 ± 4220.1 | 12.49 ± 3.97 |
| Evening (E) | 4871.0 ± 3091.9 | 10.91 ± 4.30 |
| Irregular morning (IM) | 2597.6 ± 3481.4 | 5.67 ± 5.39 |
| Irregular night (IN) | 3977.2 ± 5540.0 | 7.82 ± 6.48 |

Note: Each step-counting pattern had different daily step counts. AD had the highest, followed by M and BP, then E. IN and IM had the lowest counts. In terms of duration, the BP pattern exhibited the longest activity duration per day, followed by AD, M and E patterns. IM and IN has the lowest duration of activity.

### Clustering dominant daily step-behaviors

According to the six 24-hour step-counting patterns, we further clustered participants based on the probability of each pattern. We found five daily step-behavioral clusters; AD dominant, AD + BP, BP dominant, AD + E, and M dominant (Fig 2). Half of the participants (21) belong to the AD-dominant (S4 Table), consistent with the result that AD was the most frequent step-counting pattern. The AD+BP was the following frequent behavior. The other daily step behaviors shared minor proportions. There were no IM dominant or IN dominant step behaviors due to their minor proportions in the 24-hour step-counting patterns. Interestingly, mean values of step count amounts for each step behavior were similar (Table S4).

**Fig 2.**
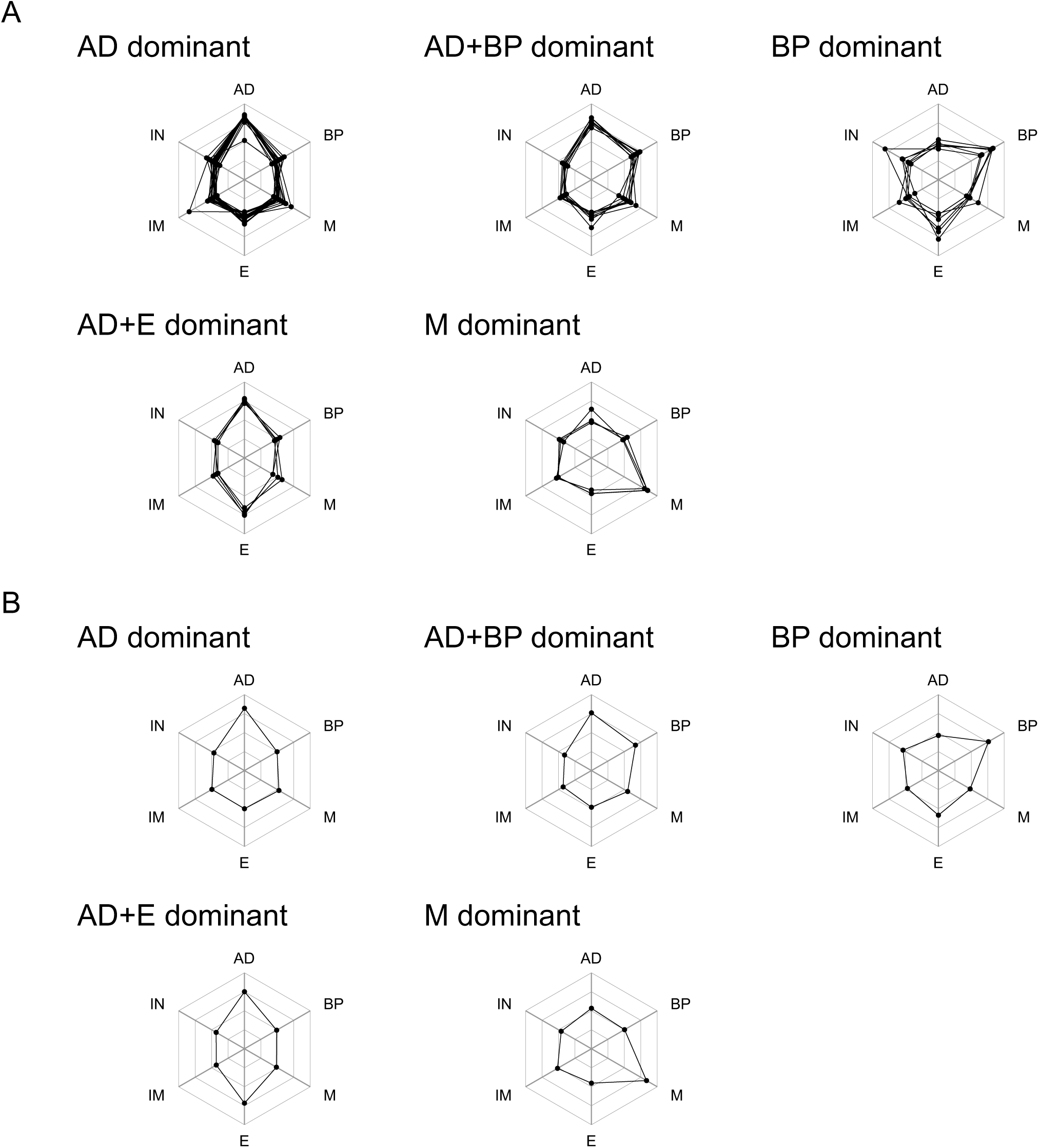
Five step behaviors. Step behavior clusters were identified using unsupervised machine learning. (A) All traces of proportion of step-counting patterns in each behavior. (B) Averaged traces (mean) of the proportion of step-counting patterns in each behavior. AD, all-day; BP, bi-phasic; M, morning; E, evening; IM, irregular morning; IN, irregular night.

Table 4 represents the probabilities of six step-counting patterns in each daily step-behavioral cluster. The AD dominant step-behavior accounts for most of the AD pattern (0.5515 ± 0.1492). AD dominant step behavior also consists of a significant portion with the M pattern (0.1227 ± 0.0888). Other step-counting patterns are less. AD+BP step-behavior is characterized by a significant share of AD and BP step-counting patterns (0.4031 ± 0.1021 and 0.2964 ± 0.0586, respectively). AD+BP step-behavior also contains a significant amount with the M step-counting pattern (0.1587±0.0748), while other step-counting patterns are less frequent. BP dominant step-behavior shares a significant proportion of the BP step-counting pattern (0.3984 ± 0.1167). This step behavior also contains a minor amount with the IN and E step-counting patterns (0.1482 ± 0.1694 and 0.2045 ± 0.1356, respectively). AD+E step-behavior consists of a similar amount with the AD and E step-counting patterns (0.4195 ± 0.0168 and 0.3815 ± 0.0771, respectively), while other step-counting patterns are less. M dominant behavior is characterized by most of the M step-counting pattern (0.6235 ± 0.1509) and also consists of less frequency of the AD and IM step-counting patterns (0.1267 ± 0.1378 and 0.1105 ± 0.0301, respectively). Though IM and IN step-counting patterns share the least number of days and do not show any dominant step-behavioral clusters, these step-counting patterns show some prevalence in all step-behavioral clusters, especially in BP dominant and M dominant step-behavioral clusters.

**Table 4.** Probability of 24-hour step-counting patterns in each daily step-behavior.

| Behavior | Mean (SD) |  |  |  |  |  |
| --- | --- | --- | --- | --- | --- | --- |
|  | AD | BP | M | E | IM | IN |
| AD dominant | 0.5515<br>(0.1492) | 0.0887<br>(0.0606) | 0.1227<br>(0.0888) | 0.0998<br>(0.0702) | 0.0886<br>(0.1190) | 0.0486<br>(0.0610) |
| AD+BP | 0.4031<br>(0.1021) | 0.2964<br>(0.0586) | 0.1587<br>(0.0748) | 0.0850<br>(0.0800) | 0.0398<br>(0.0362) | 0.0117<br>(0.0237) |
| BP dominant | 0.0740<br>(0.0468) | 0.3984<br>(0.1167) | 0.0833<br>(0.0701) | 0.2045<br>(0.1356) | 0.0915<br>(0.0752) | 0.1482<br>(0.1694) |
| AD+E | 0.4195<br>(0.0168) | 0.0916<br>(0.0422) | 0.0770<br>(0.0938) | 0.3815<br>(0.0771) | 0.0156<br>(0.0313) | 0.0147<br>(0.0294) |
| M dominant | 0.1267<br>(0.1378) | 0.0903<br>(0.0604) | 0.6235<br>(0.1509) | 0.0175<br>(0.0304) | 0.1105<br>(0.0301) | 0.0314<br>(0.0278) |

Note: This table displays probabilities of six step-counting patterns in daily step-behavioral clusters. AD dominant behavior is mostly AD pattern, with some M pattern. AD+BP is mainly AD pattern and BP pattern, with some M pattern. BP dominant behavior is primarily BP pattern, with minor IN pattern and E pattern. AD+E behavior is evenly split between AD pattern and E pattern. M dominant behavior is mostly M pattern, with some AD pattern and IM pattern. IM and IN are less common but present in all clusters, especially in M dominant and BP dominant behavior respectively.

### Scale effect evaluation by simulations

We then investigated the scale effects of day and participant numbers on cluster identification (Fig 3 and Figs S2-5). First, the number of days was randomly selected from 3 to 18 days per participant, and our procedure was applied. When 3 days of data per participant were used, only 4 step-counting patterns (Fig 3A and Fig S2A) and 3 step behaviors (Fig 3B and Fig S3A) were identified. By increasing the number of days, the number of step-counting patterns and step behaviors increased and reached a plateau over 9 days of data per participant (Fig 3A and 3B). The step-counting patterns identified from 3-18 days of data were consistent with those from the full data set (Fig 1 and Fig S2), although step behaviors were slightly different from those of the full data set (Fig 2 and Fig S3). When 15 days per participant (540 days from 36 participants) were selected, step behaviors became consistent with those from the full data set were detected set (Fig S3B).

**Fig 3.**
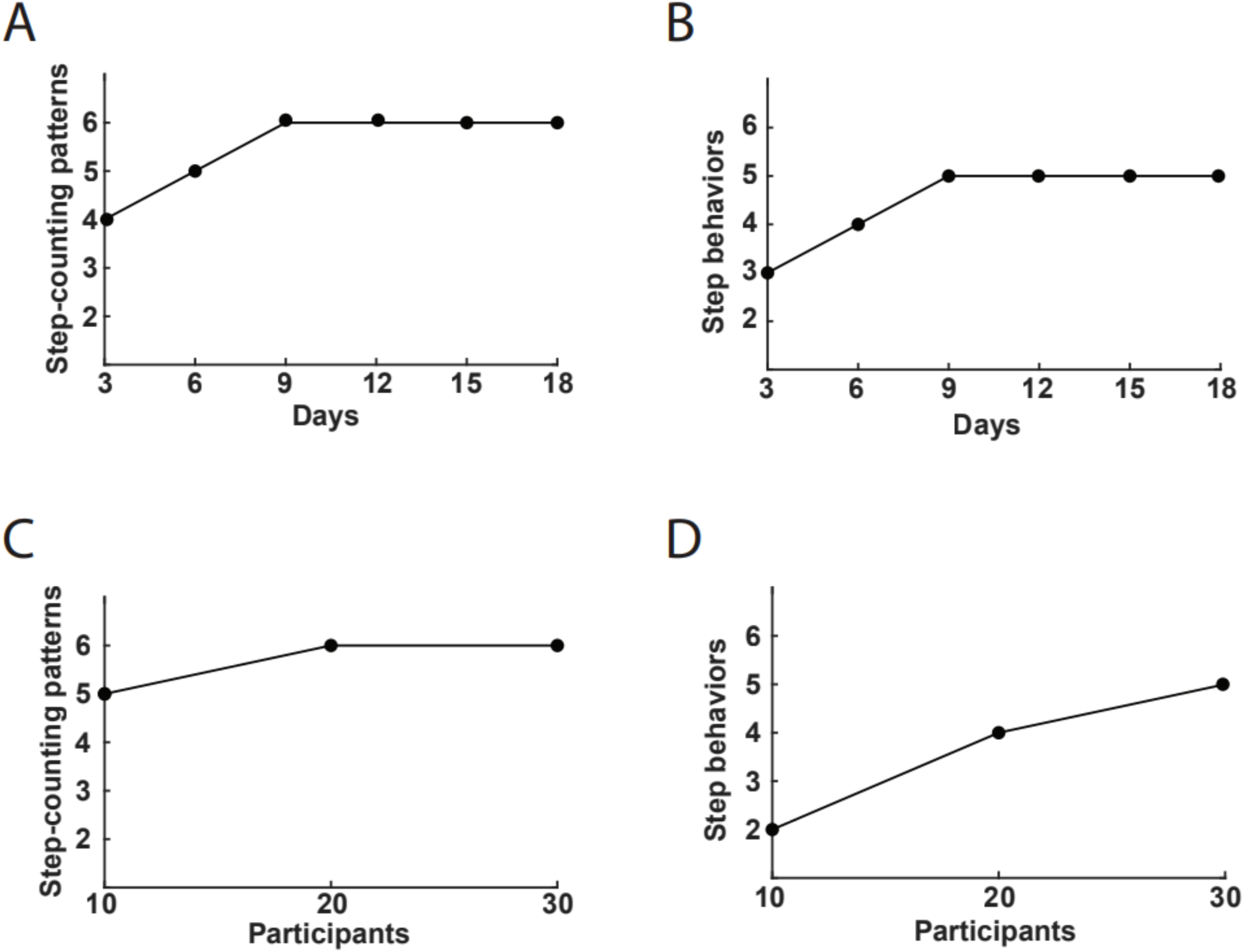
Scale effect evaluation by simulations. The effects of day and participant numbers on cluster identification were investigated. The effects of different numbers of days on identifying the numbers of step-counting patterns (A) and step behaviors (B). The effects of different numbers of participants on identifying the numbers of step-counting patterns (C) and step behaviors (D).

We then examined the scale effect of participant number on the step behavior clustering by our procedure. When 10 participants were randomly selected, 5 step-counting patterns were identified (Fig 3C and Fig S4), while only 2 step behaviors were detected (Fig 3D and Fig S5). When 20 participants, step-counting patterns increased to 6 but the number of step behaviors was four (Fig 3C and 3D). Analysis using 30 participant data set (572 days) showed a similar result to the full data set (Fig 2 and Fig S5). These results suggest that at least 15 days per participant (42 participants) or data set from 30 participants (572 days) are required to obtain robust results.

### Categorization by a traditional tertile procedure shows heterogeneity in step-counting patterns

We identified six 24-hour step-counting patterns and five daily step behavior clusters using unsupervised machine-learning approaches. Then, we compare the step amounts from these step-counting patterns to those from traditional grouping approaches such as the tertile procedure.

Table 5 summarized the composition of step-counting patterns in each tertile group. AD is the most dominant step-counting pattern in the high and mid groups (52.7% and 46.6%, respectively). However, the composition of BP, M, and E step-counting patterns differed between the high and mid groups. On the other hand, the low tertile group consists of a relatively similar proportion for each step-counting pattern. The proportion of tertile groups in each step-counting pattern was consistent with the result above (Table 6). In the AD pattern, the high tertile group is dominant (44.2%) and followed by the mid (37.9%) and low groups (17.9%). In of BP pattern, mid tertile group is the most dominant (50.7%) and followed by the high (26.0%) and low (23.3%) tertile groups. The low tertile and high tertile groups were similar (39.5%) and (38.0%) of days in the M pattern. In the E pattern, the low tertile group is the dominant (42.2%), followed by mid (31.1%) and high (21.7%). IM and IN patterns consist of low tertile groups (76.6% and 67.5%, respectively. Each group showed significantly different compositions (Table S5 – S8).

**Table 5.**
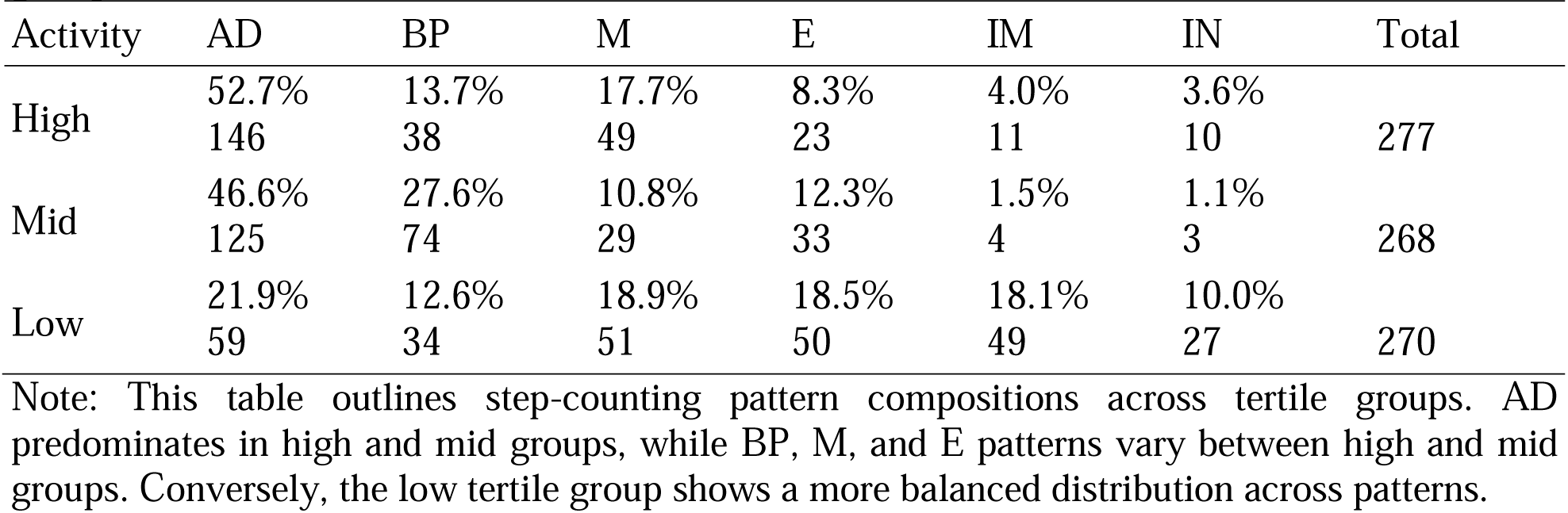
Proportion and number of days of 24-hour step-counting pattern in each tertile group.

| Activity | AD | BP | M | E | IM | IN | Total |
| --- | --- | --- | --- | --- | --- | --- | --- |
| High | 52.7%<br>146 | 13.7%<br>38 | 17.7%<br>49 | 8.3%<br>23 | 4.0%<br>11 | 3.6%<br>10 | 277 |
| Mid | 46.6%<br>125 | 27.6%<br>74 | 10.8%<br>29 | 12.3%<br>33 | 1.5%<br>4 | 1.1%<br>3 | 268 |
| Low | 21.9%<br>59 | 12.6%<br>34 | 18.9%<br>51 | 18.5%<br>50 | 18.1%<br>49 | 10.0%<br>27 | 270 |

Note: This table outlines step-counting pattern compositions across tertile groups. AD predominates in high and mid groups, while BP, M, and E patterns vary between high and mid groups. Conversely, the low tertile group shows a more balanced distribution across patterns.

**Table 6.** Proportion and number of days of tertile groups for each 24-hour step-counting pattern.

| Pattern | High | Mid | Low | Total |
| --- | --- | --- | --- | --- |
| AD | 44.2 %<br>146 | 37.9 %<br>125 | 17.9 %<br>59 | 330 |
| BP | 26.0 %<br>38 | 50.7 %<br>74 | 23.3 %<br>34 | 146 |
| M | 38.0 %<br>49 | 22.5 %<br>29 | 39.5 %<br>51 | 129 |
| E | 21.7 %<br>23 | 31.1 %<br>33 | 42.2 %<br>50 | 106 |
| IM | 17.2 %<br>11 | 6.3 %<br>4 | 76.6 %<br>49 | 64 |
| IN | 25.0 %<br>10 | 7.5 %<br>3 | 67.5 %<br>27 | 40 |

Note: Tertile group proportions align with previous findings as in Table 5. In the AD pattern, high tertile dominates, followed by mid and low. BP is mostly in the mid tertile, followed by high and low. M pattern is similar in low and high tertile. E pattern is dominated by the low tertile, followed by mid and high. IM and IN patterns are mainly in the low tertile groups.

## Discussion

In this study, we attempted to develop a procedure to cluster 24-hour step-counting patterns, as physical activity patterns, using unsupervised machine learning. We identified six step-counting patterns and five daily step behavior clusters. Comparing traditional tertile procedure (high, medium, and low) using daily step amounts, a different proportion of six 24-hour step-counting patterns were observed in each tertile group, suggesting heterogeneity in the categorization by the traditional procedure.

The “temporal patterns of physical activity” in the general term have been paid more attention in recent decades (Aqeel et al., 2021; Creasy et al., 2021; De Baere, Lefevre, De Martelaer, Philippaerts, & Seghers, 2015; Hallman, Mathiassen, Gupta, Korshøj, & Holtermann, 2015). However, the situation is confusing since this general term includes different “activity” aspects, such as simple body movements or steps (Aqeel et al., 2021), active/sedentary behaviors (Creasy et al., 2021), social behaviors (De Baere et al., 2015), or those combinations (Hallman et al., 2015). Only several studies focused on the 24-hour temporal physical activity patterns, such as body movements or steps in a narrower criterion (Aqeel et al., 2021; Niemelä et al., 2019; Smagula et al., 2015, 2018, 2022). For clustering, some previous studies extracted and used several features, such as rhythm height, timing and robustness of physical activity (Smagula et al., 2015), several parameters affecting phase shape and length of rest-activity rhythm (Smagula et al., 2018, 2022), or timing, intensity, and duration of activity (Aqeel et al., 2021). Another study used 7-day-continuous data of physical activity counts for clustering and then analyzed intensity and temporal patterns of physical activity for 24 hours (Niemelä et al., 2019). These studies adopted a two-step strategy, that is, 1) clustering by extracted features and 2) analyzing 24-hour patterns of physical activity. By contrast, our procedure first directly identified the 24-hour patterns of physical activity from bulk data of daily step-counting records using time-series clustering analysis (Fig 1) and further clustered participants depending on the probability of each pattern (Fig 2). Therefore, our strategy appears more straightforward, direct, and intuitive than previous studies.

The scale effect analyses by simulation using different numbers of randomly selected data sets indicated the limitation of our procedure when applying for a smaller number of data (Fig 3 and Figs S2-5). The analyses indicated that at least 15 days per participant (540 days from 36 participants) or data set from 30 participants (572 days) are required to obtain robust results, suggesting that over 500 days of data set could be essential. The relationship between the number of days per participant and participants remains to be investigated using a larger data set.

Conventional quantile procedure has been employed on the total amount of activity to categorize data into some clusters because of its simple application. Recently, however, quantile procedure has been pointed for a few potential problems; (1) reduction of detection power, (2) multiple comparison testing, (3) assuming the homogeneity of risk within the group, and (4) difficulty in comparing results within studies (Bennette & Vickers, 2012; Greenland, 1995). Unfortunately, these problems remain to be unsolved to date as discussed in a recent review (Jones & Ekelund, 2019). Our procedure described in this study might provide a complementary strategy to compensate for the heterogeneity in the quantile-based clustering. Our study showed an association between step-counting patterns and step amounts and heterogeneity in step-counting patterns among tertile groups based on the step amount, producing a typical example of the second problem (Tables 6 and 7). Our approach may be helpful for understanding and handling the heterogeneity derived from different physical activity patterns between/within the groups, considering it as a confounding factor.

A significant limitation of the current study is that our dataset was small and collected from office workers in the Miyagi area in the Tohoku region, Japan. As daily activity depends on job type and the geographical area where participants live, it is still challenging to mention how our procedure can be generalized for different populations. A more extensive and diverse dataset needs to be analyzed to confirm our procedure’s reliability, robustness, and reproducibility. Our approach using unsupervised machine learning to daily activity behavior and patterns may help to understand people’s prominent behaviors considering heterogeneity. Although the relationship between our results and health outcome is beyond the goal of this study, our procedure might be reliable for identifying specific patterns of physical activity and activity behavior associated with positive health outcomes. It could help the development of targeted interventions aimed at promoting healthy behavior.

## Conclusion

In our study, we introduced a new approach using unsupervised machine learning to group 24-hour step-counting patterns. This approach revealed six distinct patterns and five clusters of daily step behaviors. We found significant heterogeneity in daily step behavior categorization when applying a traditional tertile procedure based on total step amount, emphasizing the importance of considering temporal variations in activity.

## Author contribution

SSN and HI for conceptualization, methodology, data analysis, interpretation, and original draft writing; HM for writing, review, and editing; RN for supervision. All authors read and approved the final manuscript.

## Data availability

Data is available on request from the authors.

## Acknowledgments

We would like to thank all the participants and staff who have contributed to the experiment. A pioneering research support grant from Tohoku University partially supported this project. The funding body has no role in the study design, data analysis, or manuscript submission.

