## Supplementary Tables S1-S8 for "Clustering temporal step-counting patterns for 24 hours with machine learning revealed potential heterogeneity in the categorization by a traditional tertile procedure"

| **S1 Table. P values for difference of proportion between step-counting patterns (Pearson Chi-square test)** | | | | | | |
| --- | --- | --- | --- | --- | --- | --- |
| Patterns | AD | BP | M | E | IM | IN |
| AD | - |  |  |  |  |  |
| BP | 0.0008 | - |  |  |  |  |
| M | <.0001 | <.0001 | - |  |  |  |
| E | <.0001 | 0.0003 | 0.0238 | - |  |  |
| IM | <.0001 | <.0001 | <.0001 | 0.0001 | - |  |
| IN | <.0001 | <.0001 | 0.0059 | 0.0111 | 0.5834 | - |

Note: All pattern proportions exhibited significant differences except for IM and IN, which showed no significant distinction. P<0.05 is considered as significant.

| **S2 Table. P values for difference of step count amount between step-counting patterns (Tukey HSD)** | | | | | | |
| --- | --- | --- | --- | --- | --- | --- |
| Patterns | AD | BP | M | E | IM | IN |
| AD | - |  |  |  |  |  |
| BP | 0.1610 | - |  |  |  |  |
| M | 0.6743 | 0.9831 | - |  |  |  |
| E | <0.0001 | 0.0454 | 0.0078 | - |  |  |
| IM | <0.0001 | <0.0001 | <0.0001 | 0.0008 | - |  |
| IN | <0.0001 | 0.0070 | 0.0015 | 0.7515 | 0.3836 | - |

Note: AD, BP, and M patterns showed significant differences from E, IM, and IN. Notably, among E, IM, and IN patterns, a significant difference was observed specifically between E and IN. P<0.05 is considered as significant.

| **S3 Table. P values for difference of duration between step-counting patterns (Tukey HSD)** | | | | | | |
| --- | --- | --- | --- | --- | --- | --- |
| Patterns | AD | BP | M | E | IM | IN |
| AD | - |  |  |  |  |  |
| BP | 0.9816 | - |  |  |  |  |
| M | <0.0001 | <0.0001 | - |  |  |  |
| E | <0.0001 | <0.0001 | 0.0095 | - |  |  |
| IM | <0.0001 | <0.0001 | <0.0001 | <0.0001 | - |  |
| IN | <0.0001 | <0.0001 | <0.0001 | <0.0001 | 0.0313 | - |

Note: All pairs showed significant differences in duration except for AD and BP. P<0.05 is considered as significant.

| **S4 Table. Number of participants and step count amount for each daily step behaviours** | | |
| --- | --- | --- |
| Behaviour | No of Participants | Step count amount (mean ± SD) |
| AD dominant | 21 | 5773.9 ± 1346.8 |
| AD+BP | 8 | 5934.1 ± 1570.7 |
| BP dominant | 6 | 5724.3 ± 1412.8 |
| AD+E | 4 | 6722.6 ± 2144.6 |
| M dominant | 3 | 6797.8 ± 2968.5 |
| Total | 42 |  |

Note: Half of the participants exhibited AD-dominant behavior, aligning with AD being the most frequent step-counting pattern. AD+BP is the next most common behavior, while other patterns had minor representation. No participants showed dominance in IM or IN due to their low proportions in the 24-hour step-counting patterns. Remarkably, mean step count values were similar across all step behaviors.

| **S5 Table. P values for difference of proportion of step-conuting patterns in high activity group (Pearson Chi-square test)** | | | | | | |
| --- | --- | --- | --- | --- | --- | --- |
| Patterns | AD | BP | M | E | IM | IN |
| AD | - |  |  |  |  |  |
| BP | 0.0002 | - |  |  |  |  |
| M | 0.2228 | 0.0334 | - |  |  |  |
| E | <0.0001 | 0.4283 | 0.0070 | - |  |  |
| IM | 0.0001 | 0.1633 | 0.0033 | 0.4762 | - |  |
| IN | 0.0199 | 0.8953 | 0.1323 | 0.6705 | 0.3343 | - |

Note: In the high activity group, AD significantly differs from all patterns except M. BP only differs significantly from M. M differs significantly from E and IM. E does not differ significantly from IM and IN. IM does not differ significantly from IN. p<0.05 is considered as significance.

| **S6 Table. P values for difference of proportion of step-counting patterns in medium activity group (Pearson Chi-square test)** | | | | | | |
| --- | --- | --- | --- | --- | --- | --- |
| Patterns | AD | BP | M | E | IM | IN |
| AD | - |  |  |  |  |  |
| BP | 0.0090 | - |  |  |  |  |
| M | 0.0017 | <0.0001 | - |  |  |  |
| E | 0.2087 | 0.0019 | 0.1343 | - |  |  |
| IM | <0.0001 | <0.0001 | 0.0048 | 0.0001 | - |  |
| IN | 0.0001 | <0.0001 | 0.0346 | 0.0031 | 0.8045 | - |

Note: In the medium activity group, AD significantly differs from all patterns except E. BP shows significant differences with all patterns. M does not differ significantly from E. E differs significantly from IM and IN. IM does not differ significantly from IN. p<0.05 is used for significance.

| **S7 Table. P values for difference of proportion of step-counting patterns in low activity group (Pearson Chi-square test)** | | | | | | | | | |
| --- | --- | --- | --- | --- | --- | --- | --- | --- | --- |
| Patterns | AD | | BP | M | | E | IM | | IN |
| AD | - | |  |  | |  |  | |  |
| BP | 0.1699 | | - |  | |  |  | |  |
| M | <0.0001 | | 0.0036 | - | |  |  | |  |
| E | <0.0001 | | 0.0001 | 0.2394 | | - |  | |  |
| IM | <0.0001 | | <0.0001 | <0.0001 | | 0.0002 | - | |  |
| IN | <0.0001 | | <0.0001 | 0.0019 | | 0.0282 | 0.3107 | | - |
| Note: In the low activity group, AD and BP patterns are similar but differ significantly from others. M and E patterns are alike but significantly differ from all others. IM significantly differs from all except IN. Significance is considered as p<0.05.  S8 Table. P values for difference of proportion of step-counting patterns between tertile groups (Pearson Chi-square test) | | | | | | | | | |
| Pattern | | Low-Mid | | | Low-High | | | High-Mid | |
| AD | | <0.0001 | | | <0.0001 | | | 0.1568 | |
| BP | | <0.0001 | | | 0.6970 | | | 0.0001 | |
| M | | 0.0085 | | | 0.7167 | | | 0.0221 | |
| E | | 0.0463 | | | 0.0004 | | | 0.1232 | |
| IM | | <0.0001 | | | <0.0001 | | | 0.0770 | |
| IN | | <0.0001 | | | 0.0029 | | | 0.0568 | |

Note: Within the tertile groups, AD pattern is significant across all except high-mid active groups. BP pattern is significant across all except low-high group, as is M pattern. E does not show significance within high-mid active groups. IM is significant across all except high-mid group, while IN shows significance across all groups.
