## Supplementary figure S1-S5 for "Clustering temporal step-counting patterns for 24 hours with machine learning revealed potential heterogeneity in the categorization by a traditional tertile procedure"

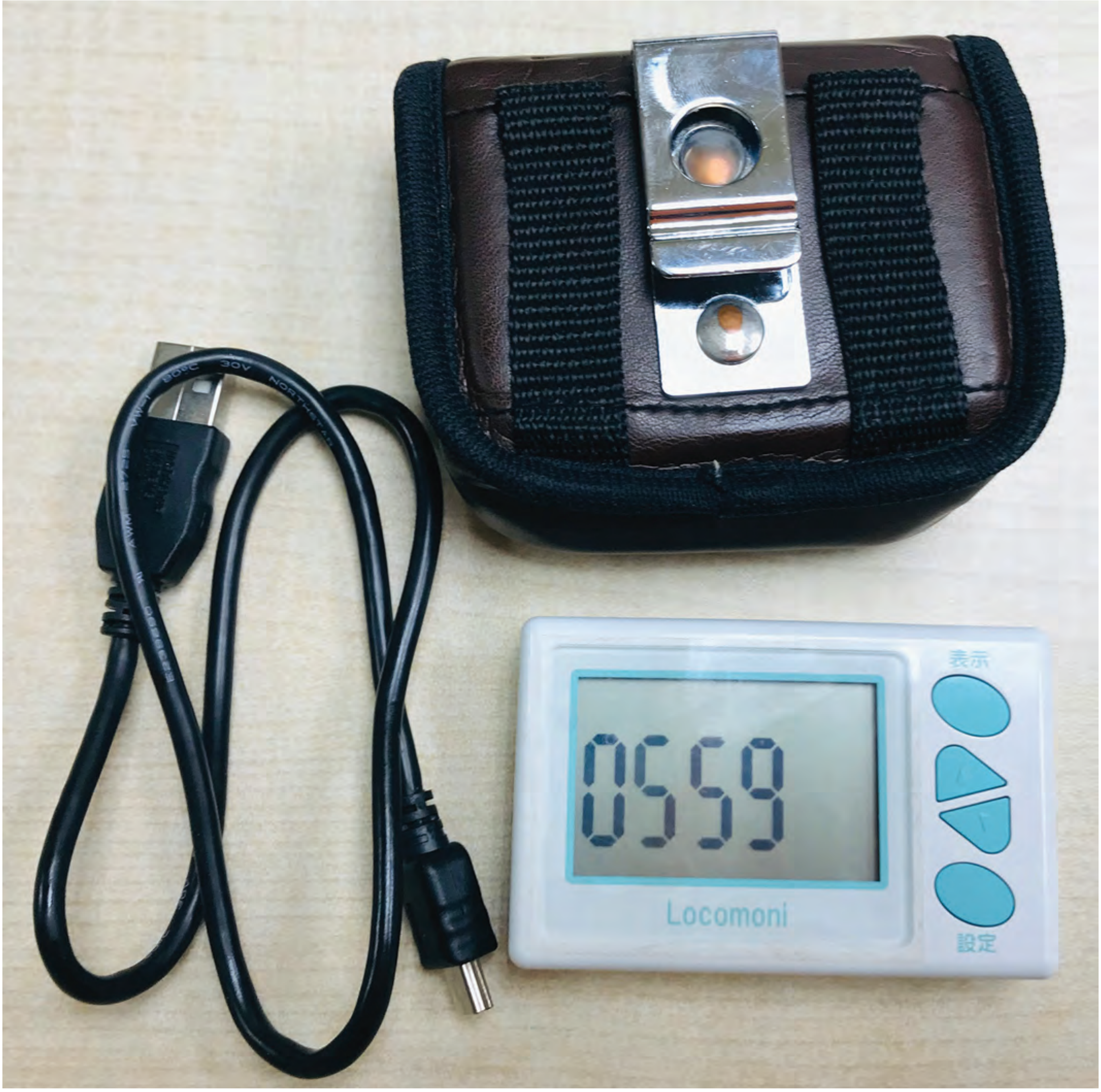


Fig. S1. Image of the hip-worn triaxial accelerometer.

Supplementary Figure 1

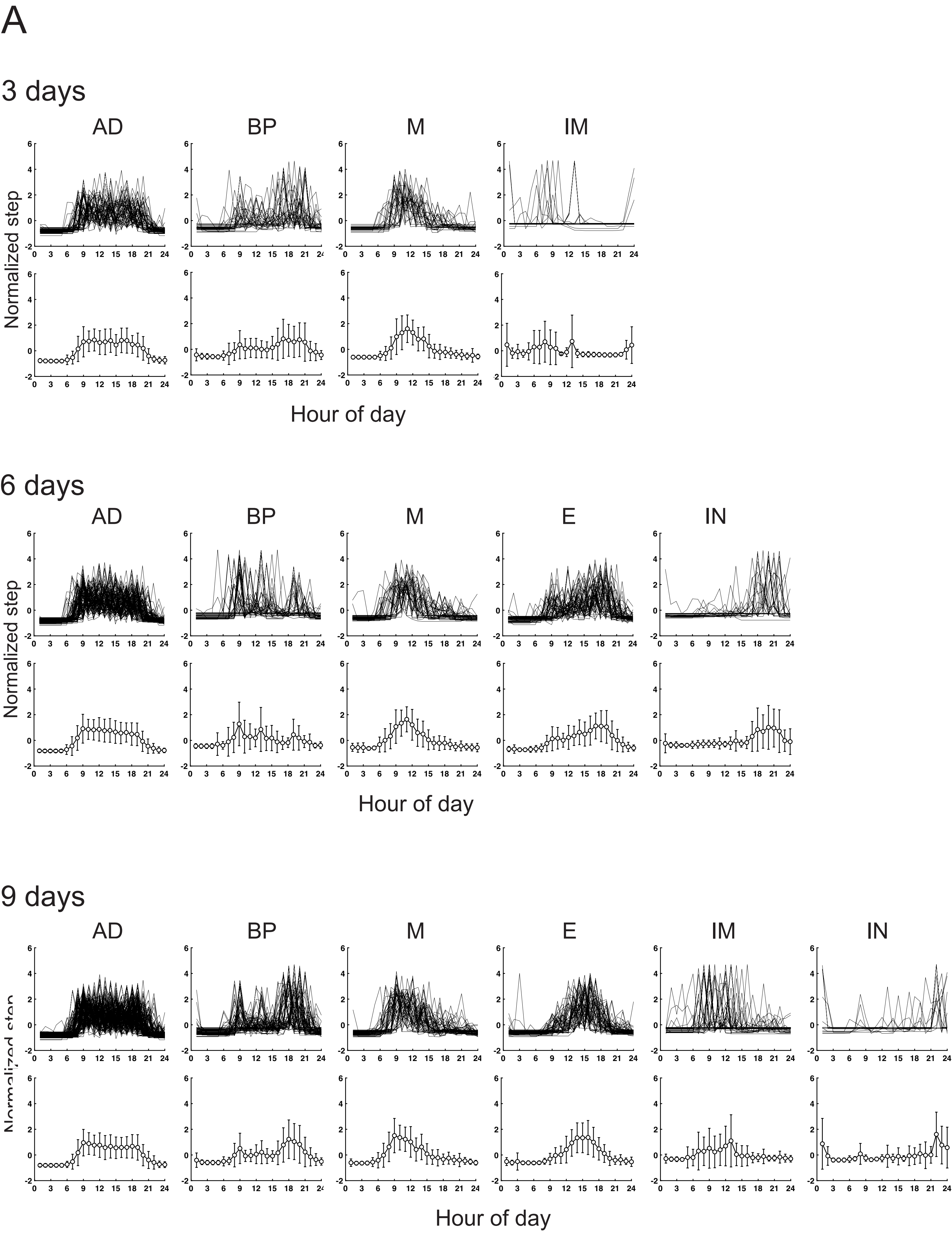


Supplementary Figure 2

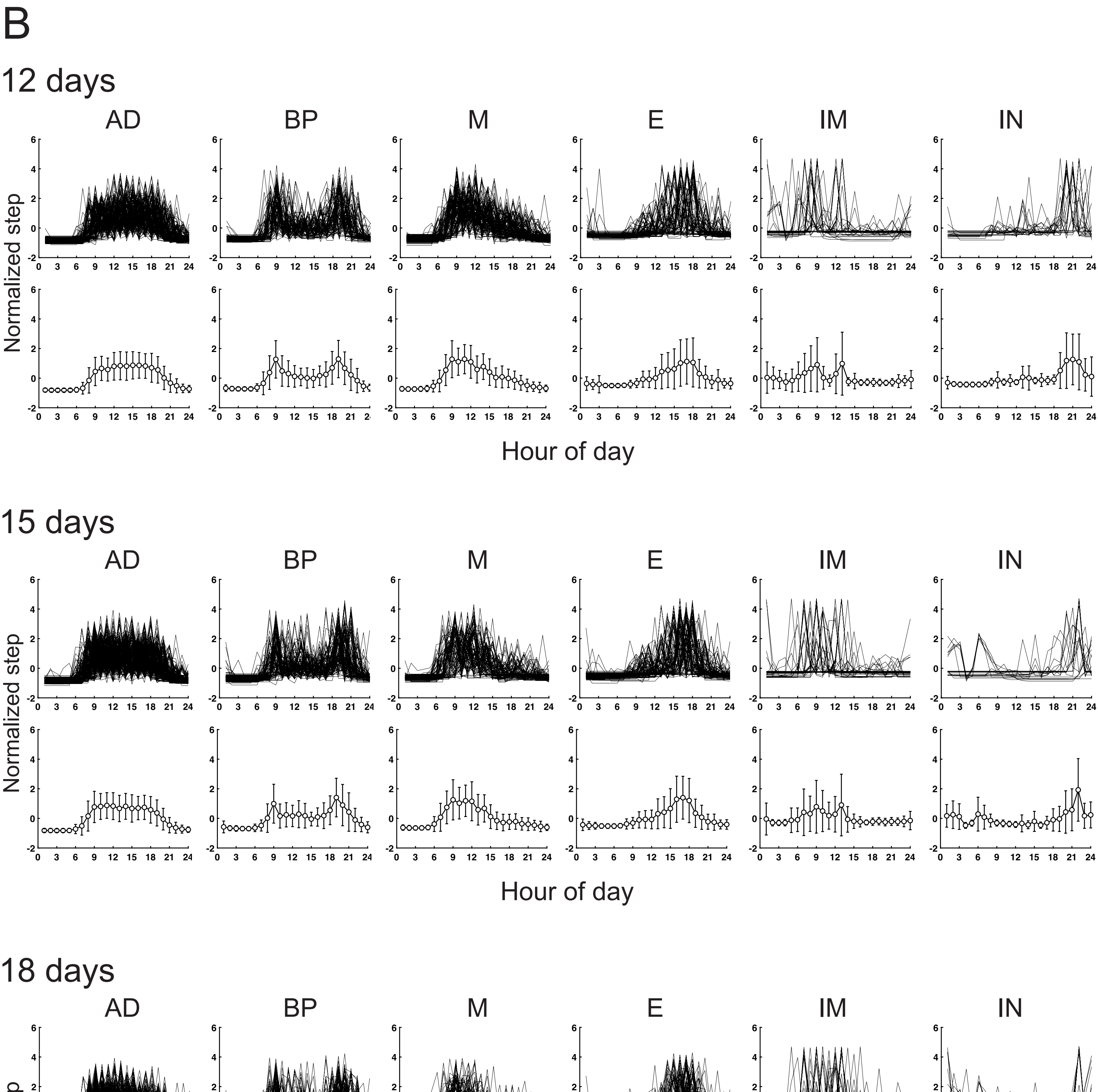


Fig. S2. Scale effect simulation for step-counting patterns on the number of days. Temporal step-counting pattern for 24 hours identified using unsupervised machine learning. (A) All traces and their respective Averaged traces (mean ± SD) of step-counting activity of each pattern for 3 days, 6 days and 9 days (B) All traces and their respective Averaged traces (mean ± SD) of step-counting activity of each pattern for 12 days, 15 days and 18 days. AD, all-day; BP, bi-phasic; M, morning; E, evening; IM, irregular morning; IN, irregular night.

Supplementary Figure 2

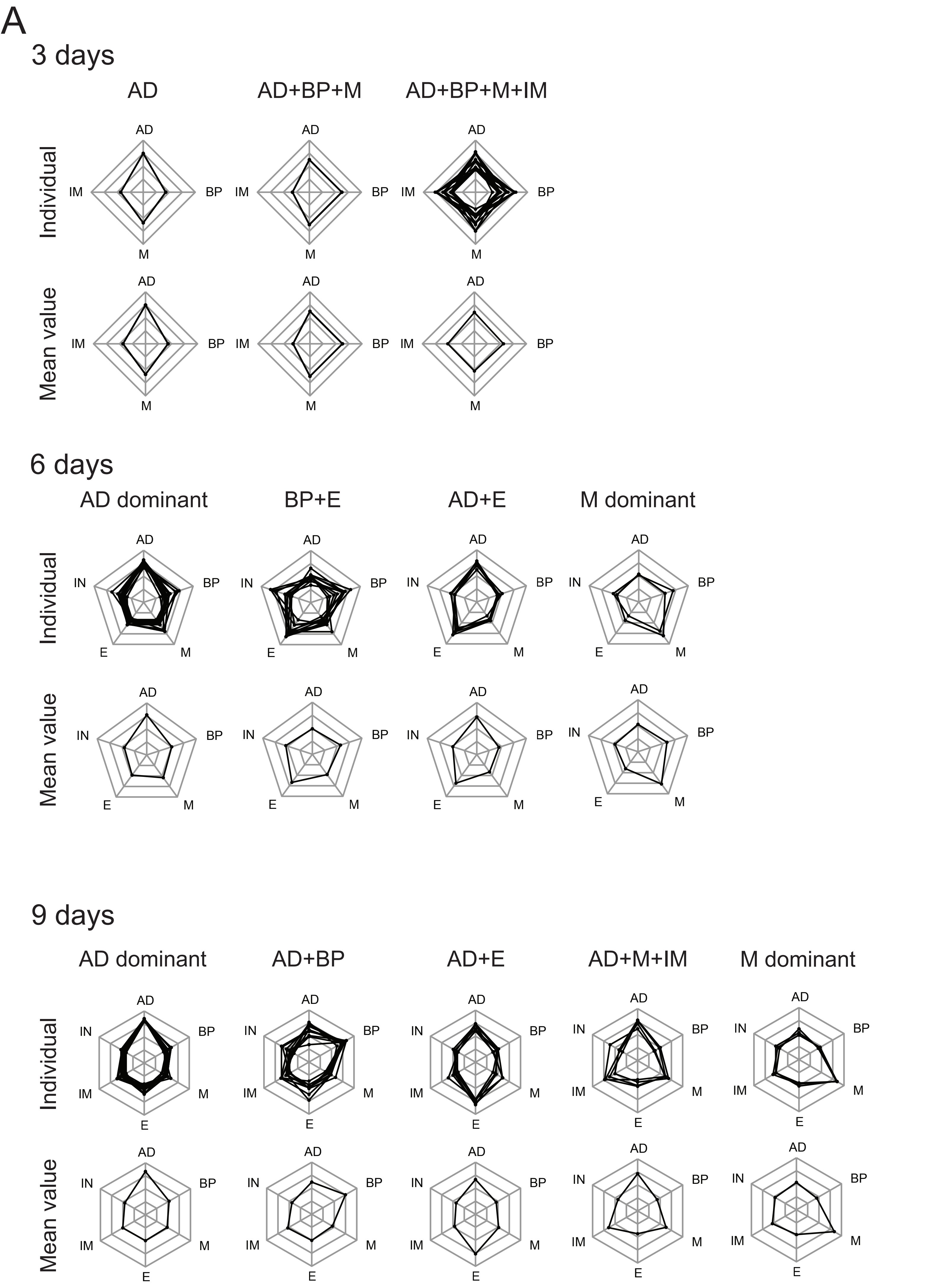


Supplementary Figure 3

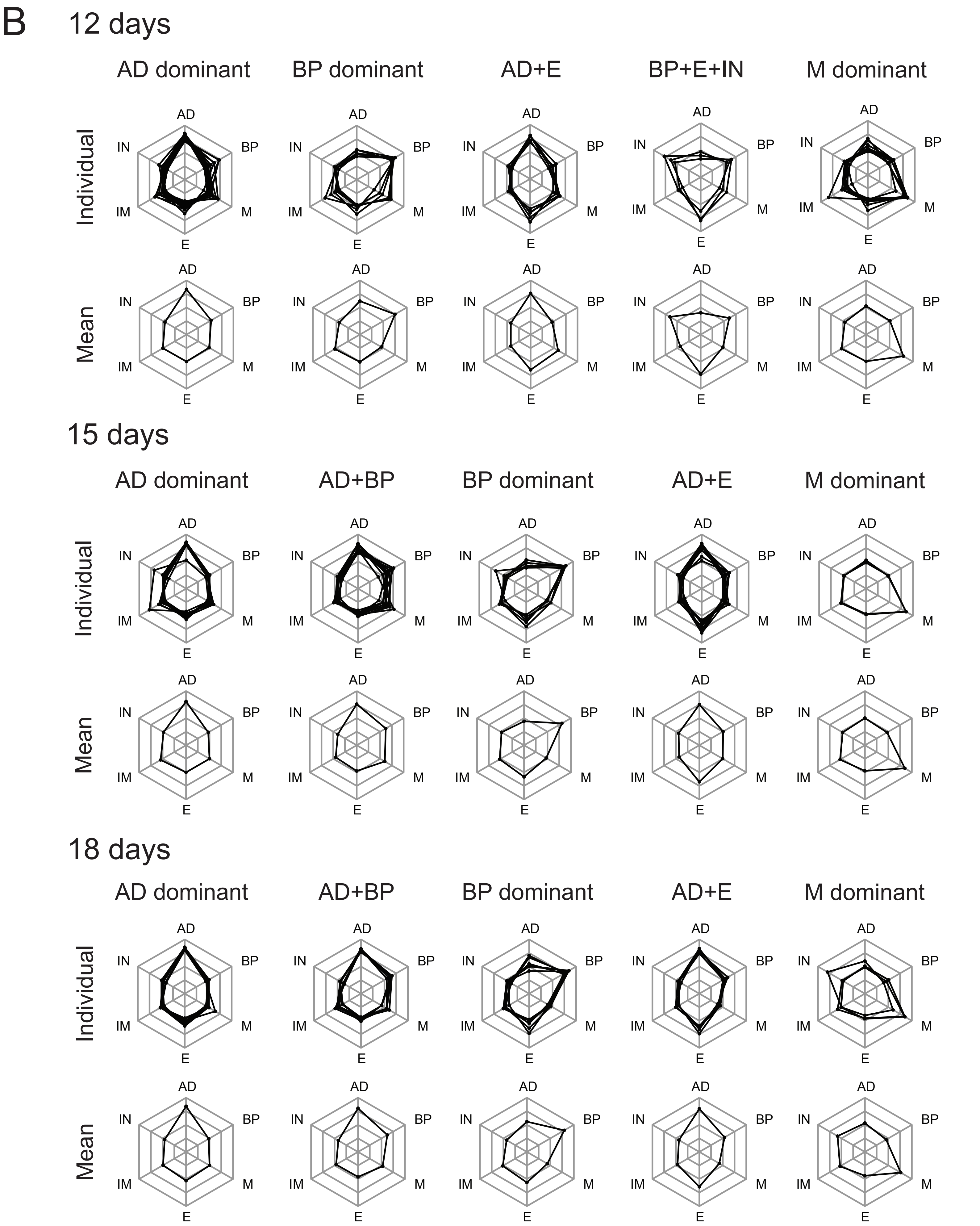


Fig. S3. Scale effect simulation for step behaviors on the number of days. Step behavior clusters were identified using unsupervised machine learning. (A) All traces of proportion and their respective averaged traces (mean) of the proportion of step-counting patterns in each behavior for 3 days, 6 days and 9 days. (B) All traces of proportion and their respective averaged traces (mean) of the proportion of step-counting patterns in each behavior for 12 days, 15 days and 18 days. AD, all-day; BP, bi-phasic; M, morning; E, evening; IM, irregular morning; IN, irregular night.

Supplementary Figure 3

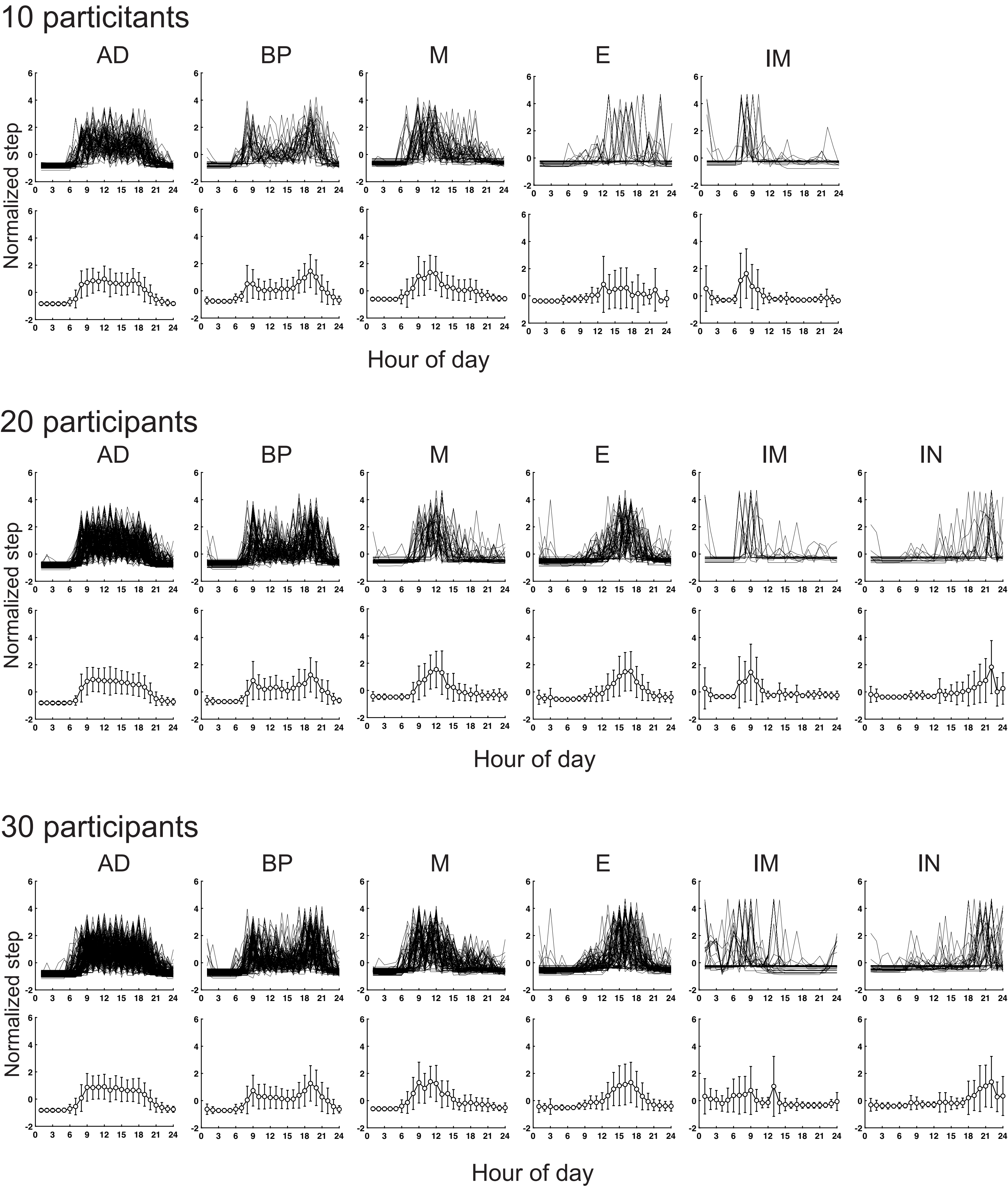


Fig. S4. Scale effect simulation for step-counting patterns on the number of participants.Temporal step-counting pattern for 24 hours identified using unsupervised machine learning. All traces and their respective Averaged traces (mean ± SD) of step-counting activity of each pattern for 10 participants, 20 participants and 30 participants. AD, all-day; BP, bi-phasic; M, morning; E, evening; IM, irregular morning; IN, irregular night.

Supplementary Figure 4

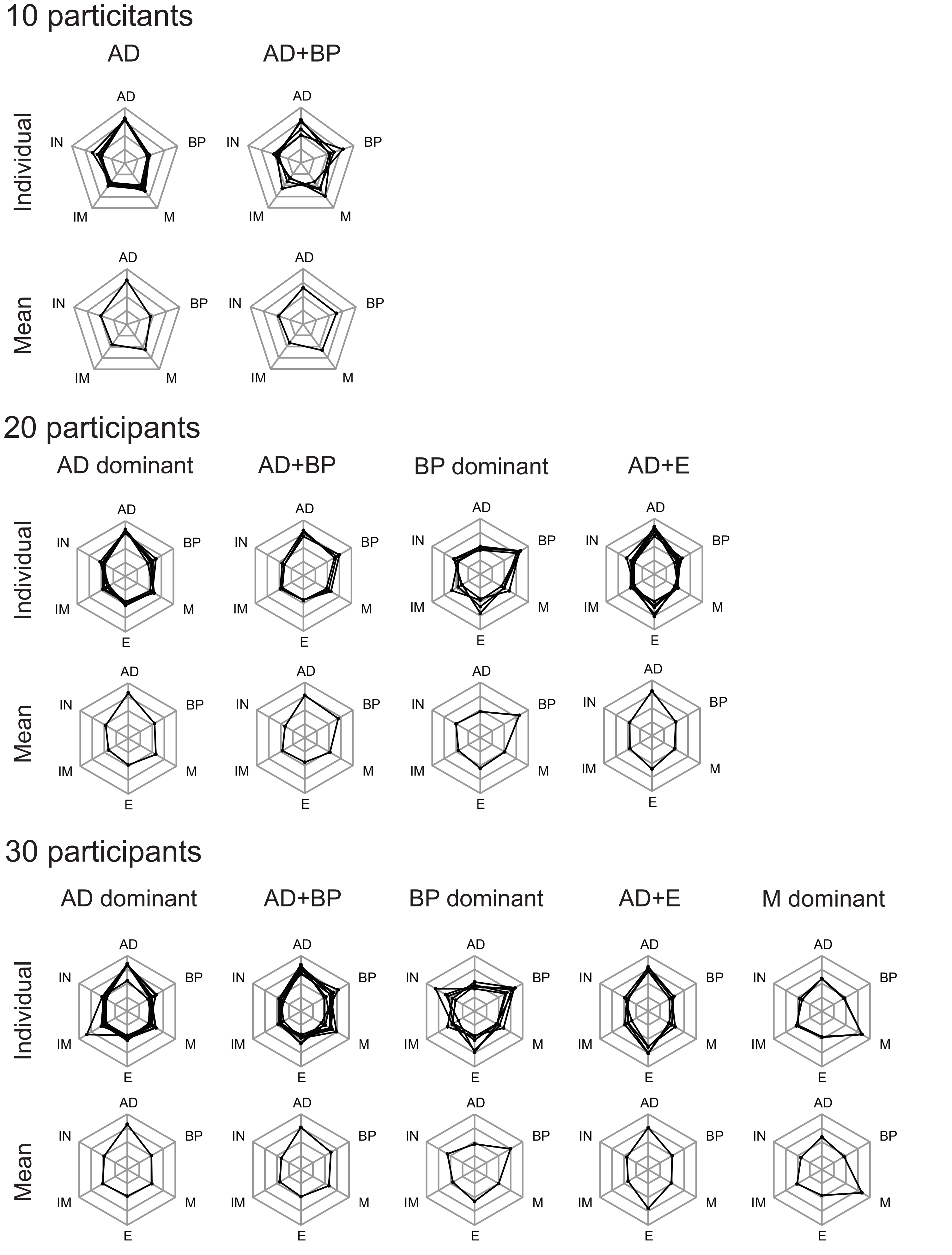


Fig. S5. Scale effect simulation for step behaviors on the number of participants. Step behavior clusters were identified using unsupervised machine learning. All traces of proportion and their respective averaged traces (mean) of the proportion of step-counting patterns in each behavior for 10 participants, 20 participants and 30 participants. AD, all-day; BP, bi-phasic; M, morning; E, evening; IM, irregular morning; IN, irregular night.

Supplementary Figure 5
